# N-Acetyl Cysteine Alleviates Coxsackievirus B-Induced Myocarditis by Suppressing caspase-1

**DOI:** 10.1101/732677

**Authors:** Yao Wang, Shuoxuan Zhao, Yang Chen, Ying Wang, Tianying Wang, Xiaoman Wo, Yanyan Dong, Jian Zhang, Weizhen Xu, Cong Qu, Xiaofeng Feng, Xiaoyu Wu, Yan Wang, Zhaohua Zhong, Wanran Zhao

## Abstract

Viral myocarditis caused by Coxsackievirus B (CVB) infection is a severe inflammatory disease of the myocardium, which may develop to cardiomyopathy and heart failure. No effective medicine is available to treat CVB infection. Here we evaluated the anti-CVB effect of N-acetyl cysteine (NAC), a widely used antioxidant. NAC significantly alleviated myocarditis and improved the overall condition of CVB type 3 (CVB3)-infected mice. Importantly, NAC treatment suppressed viral replication in both myocardium and cell culture. We show that NAC inhibited CVB3 replication when it was applied at the early stage of CVB3 infection. NAC’s antiviral mechanism, while independent of its antioxidant property, relies on its inhibition on caspase-1 activation, since the knockdown of caspase-1 blocked CVB3 replication. Moreover, NAC promotes procaspase-1 degradation via ubiquitin proteasome system, which may further contribute to the inhibited activity of caspase-1. NAC also inhibits the activity of viral proteases. Taken together, this study shows that NAC exerts potent anti-CVB effect by inhibiting caspase-1 and viral proteases. This study suggests that NAC can be a safe therapeutic option for CVB-induced myocarditis.

## Introduction

Myocarditis is defined as the inflammation of myocardium, which can be caused by the infection of various pathogens ^1, 2^. Viral myocarditis is a common disease which represents the significant cause of cardiac death, especially among young adults ^2-4^. Coxsackievirus B (CVB), a single-stranded, positive-sensed RNA virus in the *Enterovirus* genus of *Picornaviridae* family, is the leading pathogen that causes myocarditis ^5, 6^. Although acute viral myocarditis is often subclinical and the recovery may be spontaneous, severe myocarditis with the necrosis of myocardium and fatal ventricular tachycardia has been observed ^7, 8^. Increasing evidence has suggested that a substantial number of patients with viral myocarditis progress to dilated cardiomyopathy, which may lead to heart failure and cardiac death ^9^. The pathogenesis of CVB infection has not been completely understood ^5, 10^. Treatment to viral myocarditis relies primarily on supportive care due to the lack of specific antiviral agents.

CVB includes six serotypes (CVB1 – CVB6). The genome of CVB is a single-stranded, positive-sensed RNA about 7.4 kb, which encodes four capsid proteins (VP1 - VP4) and six nonstructural proteins (2A, 2B, 2C, 3A, 3C, and 3D). 3D is the RNA-dependent RNA polymerase (3D^pol^) which is responsible for the replication of viral genome, while 2A and 3C are viral proteases (2A^pro^ and 3C^pro^) which trans-cleave the viral polyprotein and are required for the maturation and assembly of viral particles ^5^. Studies have shown that 2A^pro^ and 3C^pro^ also cleave and degrade cellular proteins involved in translation and innate immunity such eukaryotic initiation factor 4G (eIF4G), mitochondrial antiviral-signaling protein (MAVS), and retinoic acid inducible gene I (RIG-I) ^11, 12^. Through cleaving eIF4G, a key component in the pre-translation initiation complex, CVB shuts down the cap-dependent translation and hence blocks cellular protein synthesis ^13^. Viral translation, however, is not affected, because picornaviral RNA is uncapped and the translation initiation uses an internal ribosome entry site (IRES) in the 5’-untranslated region (5’-UTR) ^14, 15^

To benefit viral replication, CVB also manipulates other cellular machineries such as ubiquitin proteasome system (UPS) ^16, 17^. UPS is an important non-lysosomal protein degradation mechanism for eukaryotic cells, and it is involved in fundamental cellular processes ^18^. Proteins targeted for destruction by UPS are often short-lived regulatory proteins such as cyclins, p53, and p57 ^18, 19^. In addition, UPS also targets misfolded proteins with abnormal configurations which could be harmful to the cell. Upon ubiquitination, a chain of multiple copies of ubiquitin (UB) is linked to the substrate protein, which is then degraded by 26S proteasome ^18-20^. It has been demonstrated that UPS is utilized during CVB3 infection, while dysregulated UPS showed negative impact on CVB3 replication ^16, 17^.

CVB infection typically causes inflammatory injury of the myocardium characterized by the damage of cardiac muscles and the infiltration of inflammatory cells ^1, 21^. The rapid production of proinflammatory cytokines has been seen in both patients and CVB-infected mice shortly after the infection of CVB ^22, 23^. These cytokines include IL-6, IL-1β, and TNF-α ^23, 24^. Enhanced production of inflammatory cytokines and chemokines is associated with severe impaired cardiac function and myocardial fibrosis ^25^. Proinflammatory cytokines also lead to aberrant mitochondrial metabolism of the cardiomyocytes, which further contributes to the dysfunction of the CVB-infected heart ^26^. The maturation and release of the proinflammatory cytokines IL-1β and IL-18 require active caspase-1, which proteolytically cleaves the precursors of IL-1β and IL-18 ^27^.

Caspase-1, one of the aspartic acid-specific cysteine proteases, is best known for its role as the inflammation initiator ^28, 29^. Caspase-1 is synthesized as zymogen and becomes active when recruited into inflammasomes in response to various pathogen-associated molecular patterns (MAMP) or danger-associated molecular patterns (DAMP) ^30^. Once recruited into the inflammasome, caspase-1 undergoes autoproteolysis to form the active heterodimer composed of caspase-1 p20 and p10 ^27, 28^. Active caspase-1 cleaves the precursors of IL-1β, IL-18, and IL-33, which then released from the cell to trigger inflammation ^27^. The generation of pro-inflammatory cytokines represents the first line of the defense mechanism against the invasion of pathogens ^31, 32^. Excessive caspase-1 activation, however, is involved in various pathological conditions such as cardiovascular disease ^33^, chronic kidney disease ^34^, and inflammatory bowel disease ^35^.

In addition to triggering pyroptosis and inflammation, caspase-1 might be involved in the pathogenesis of viral infection. During the infection of herpes simplex virus and vaccinia virus, caspase-1 suppresses the stimulator of interferon genes (STING)-mediated production of interferon (IFN) ^36^. Zika virus evades the host immune response through stabilizing caspase-1 by its nonstructural protein NS1, leading to the cleavage of cyclic guanosine monophosphate-adenosine monophosphate (cGAMP) synthase (cGAS) ^37^. Caspase-1 induces cell death contributes to the depletion of CD4^+^ T cells in human immunodeficiency virus (HIV)-infected patients ^38^. We previously demonstrated that CVB infection induced caspase-1 activation. surprisingly, we found that inhibition of caspase-1 with specific inhibitors not only alleviated the myocardial inflammation but also significantly inhibited viral replication, suggesting a novel therapeutic strategy against CVB-induced myocarditis ^39^.

N-acetyl cysteine (NAC), an amino acid derivative, is among the list of essential medicines issued by the World Health Organization ^40^. As the precursor of glutathione, an important cellular antioxidant molecule, NAC has been used for the treatment of various conditions with oxidative stress such as paracetamol overdose ^41, 42^, traumatic injury ^43^, as well as liver and renal failure ^44, 45^. *In vitro* study has demonstrated that the addition of NAC to cultured cells suppresses the replication of EV71 ^46^. NAC could also alleviate the liver injury caused by Dengue virus infection by promoting the production of interferon 47. So far there is no report available on the role of NAC in the therapy of CVB infection.

The present study aims to investigate the potential application of NAC in the CVB type 3 (CVB3)-induced myocarditis. Interestingly, our *in vivo* and *in vitro* data reveal that NAC can simultaneously suppress CVB3 replication and inflammatory response through blocking caspase-1 and viral proteases. Given that NAC is clinically approved medicine, NAC may therefore be a promising agent for the treatment of CVB-related myocarditis.

## Results

### NAC shows cardio-protective and antiviral effect for mice infected with CVB3

NAC is an established antioxidant. Its anti-inflammatory property has also been well demonstrated ^48, 49^. Viral myocarditis caused by CVB3 infection is primarily the inflammation of myocardium ^1^. Treatment with anti-inflammatory medicines improved heart function and reduced the serum level of proinflammatory cytokine TNF-α ^50, 51^. Motivated by the anti-inflammatory feature of NAC, we treated CVB3-infected mice with NAC, anticipating that NAC would suppress myocardial inflammation.

To this end, we used CVB3-infected Balb/c mice, the typical mouse model for the study of CVB3 infection. Newborn Balb/c mice within 3- to 5-day-old were infected with 10^6^ TCID_50_ of CVB3 intraperitoneally. Mice were given 15 mg/kg (body weight) of NAC intraperitoneally twice a day for five consecutive days starting at 12 h of post-infection (p.i.). Control mice was infected with CVB3 and treated with phosphate buffered saline (PBS). At day 5 of p.i., mice were euthanized and mouse hearts were collected and subjected to histopathological observation.

As shown in Figure 1, CVB3 infection caused debilitating changes of the mice manifested by the coarse fur (Figure 1A) and the decline of body weight (Figure 1B). One third (3/9) of the mice infected with CVB3 were alive at day 5 of p.i. In contrast, CVB3-infected mice treated with NAC showed improved fur appearance and slowly increased body weight. Two third (6/9) of the CVB3-infected mice treated with NAC were still alive at day 5 of p.i. (Figure 1C).

**Figure 1.**
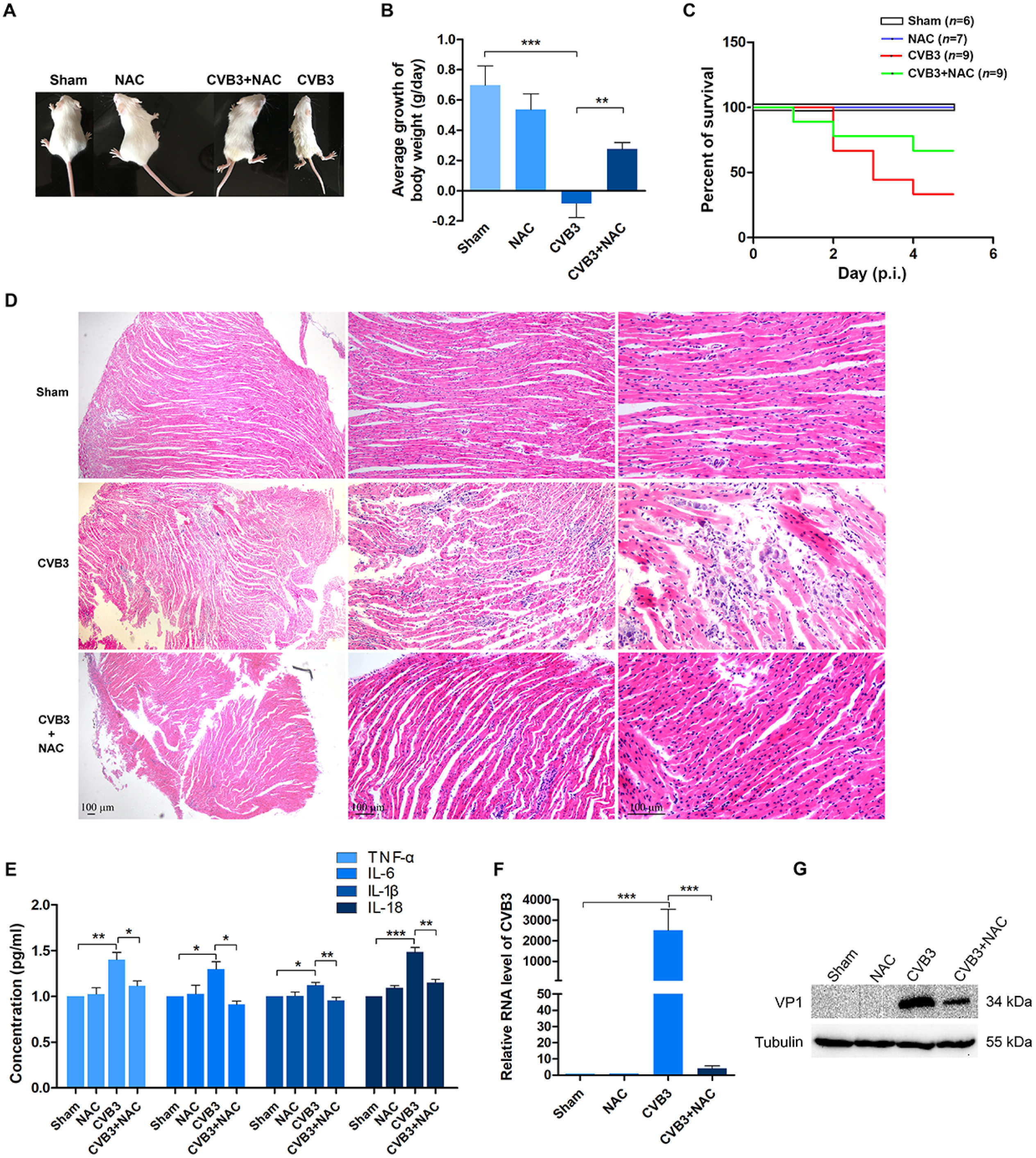
NAC exerts anti-CVB3 and cardio-protective effect. Newborn mice were infected with CVB3 (at 10^6^ TCID_50_). 15 mg/kg (body weight) of NAC was injected intraperitoneally at 12 h of post-infection twice a day for 5 days. *n* = 6-9. (A) The overall condition of the mice was shown at the end of day 5 of post-infection. (B) The average change of mouse body weight per day was calculated according to the equation: Σ (present day – previous day)/5. (C) The overall survival rate of the mice was determined at the end of day 5 of post-infection. (D) After mice were euthanized, mouse hearts were collected and subjected to histological study by HE staining. (E) Total proteins were extracted from the myocardium. Inflammatory cytokines were determined by ELISA. (F) RNA of the myocardium was extracted. CVB3 genome were determined by RT-PCR. (G) VP1 of CVB3 in the myocardium was determined by Western blotting. Experiments were repeated three times. \**P* < 0.05; * \**P* < 0.01; * * \**P* < 0.001.

Histopathological analysis shows that the infection of CVB3 resulted in significant myocardial injury represented by the damaged myofibril foci and the infiltration of inflammatory cells, while the treatment of NAC dramatically improved the morphology of myocardium with significantly mild myocardial injury and the diminished infiltration of mononuclear cells (Figure 1D), demonstrating that NAC has cardio-protective effect. The results of ELISA show that the levels of the proinflammatory cytokines including TNF-α, IL-6, IL-1β, and IL-18 were significantly reduced in the CVB3-infected myocardium with NAC treatment (Figure 1E). Collectively, these data indicate that NAC clearly inhibited cardiac inflammation resulted from CVB3 infection.

The improved morphology and inflammation of the myocardium of CVB3-infected mice with NAC treatment motivated us to ask the question how the viral load was changed. To show viral replication, total RNAs and proteins were extracted from the myocardium. Viral genome and VP1 of CVB3 were determined by RT-PCR and Western blotting, respectively. The abundance of viral genome (Figure 1F) and the level of VP1 (Figure 1G) in the myocardium infected with CVB3 were clearly decreased, indicating that viral replication is suppressed. Collectively, these data show that NAC exerts anti-inflammatory and antiviral effect for CVB3-induced myocarditis.

### NAC inhibits the replication of CVB3 in HeLa cells

We next used *in vitro* approaches to confirm the antiviral effect of NAC. HeLa cells were infected with CVB3 at MOI of 1 for 24 h. NAC was added to the culture medium at the same time as viral infection began. As shown in Figure 2, the application of NAC significantly suppressed the cytopathic effect induced by CVB3 infection (Figure 2A) and improved the viability of the cells (Figure 2B). Importantly, compared to the CVB3-infected cells without NAC treatment, viral 3D RNA polymerase (3D^pol^) (Figure 2C), viral RNA (Figure 2D), as well as the production of virions (Figure 2E) were significantly reduced in the cells treated with NAC. These results indicate that NAC exerts potent inhibition on CVB3 replication.

**Figure 2.**
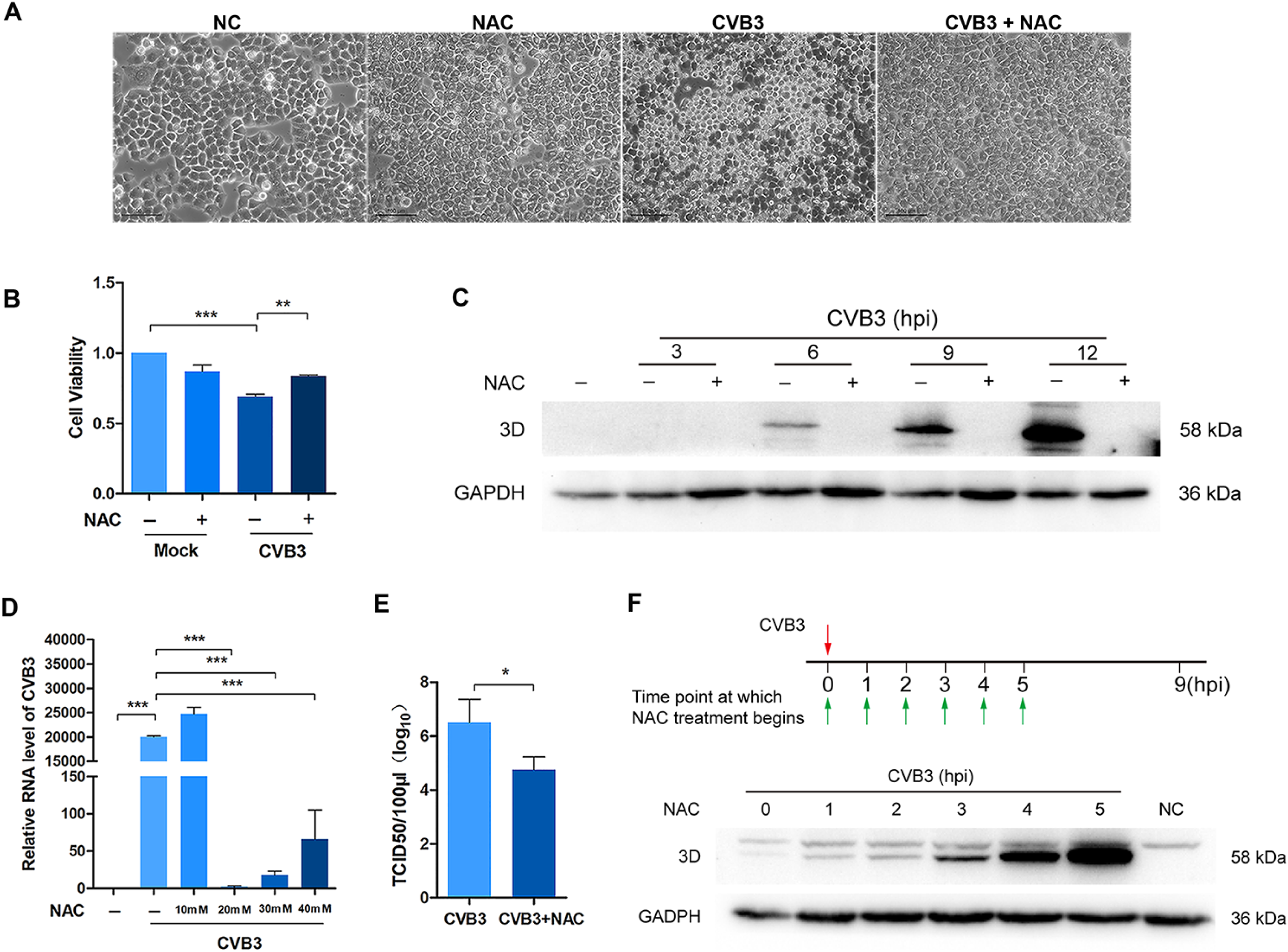
NAC inhibits the replication of CVB3 in HeLa cells. HeLa cells were infected with CVB3 (MOI = 1) for 24 h in the medium with or without the supplement of NAC (20 mM). (A) Cytopathic effect of CVB3 was viewed in microscope. (B) Cell viability was determined by Cell Counting Kit-8. (C) HeLa cells were infected with CVB3 at MOI of 1 and cultured in the medium containing NAC at 20 mM. Cells were collected at various time points of post-infection and subjected to Western blot analysis. (D) HeLa cells were infected with CVB3 at MOI of 1 and cultured in the medium supplemented with various concentration of NAC for 24 h. Total RNA was extracted and viral RNA was determined by RT-qPCR. (E) HeLa cells were infected with CVB3 at MOI of 1 for 24 h. Cells were collected and subjected to three freeze-thaw cycles. Quantification of CVB3 was performed. (F) HeLa cells were infected with CVB3. NAC (20 mM) was added to the culture medium at various time points of post-infection. Cells were collected at 9 h of post-infection and subjected to Western blot analysis. Experiments were repeated three times. *n* = 3. \**P* < 0.05; * \**P* < 0.01; * * \**P* < 0.001. hpi: hours of post-infection.

To further understand the antiviral activity of NAC during the course of CVB3 infection, HeLa cells were infected with CVB3 (MOI =1) for 9 h, and NAC was added to the culture medium at various time points of post-infection (p.i.) (Figure 1F). When NAC was added at 2 h of p.i. or earlier, the synthesis of viral 3D^pol^ was almost completely blocked. When NAC was added at 3 or 4 h of p.i., markedly reduced levels of 3D^pol^ were still observed. However, when NAC was added to the culture medium at 5 h of p.i., the synthesis of viral 3D^pol^ was not affected. These data demonstrate that NAC shows antiviral effect when it is applied at the early stage of CVB3 replication.

We found that the maximum inhibition on CVB3 replication was achieved (Figure 2D), when NAC was used at 20 mM. In the meantime, cell viability was still maintained (Figure 2B). Thus, 20 mM of NAC was used for the remaining *in vitro* experiments in this study.

### The antiviral effect of NAC is independent of its antioxidant activity

NAC is well-known as ROS scavenger, since it is believed to be the precursor of glutathione, an important antioxidant of the cell ^52^. It has been reported that certain plant RNA virus exploits the increased ROS of the host cell to promote viral replication ^53^. Moreover, inhibited generation of ROS by NAC has negative impact on the replication of EV71 in cultured cells ^46^. Thus, we asked the question whether or not the suppressed replication of CVB3 was due to the reduced generation of ROS as a result of NAC treatment.

To this end, HeLa cells were cultured in the medium containing NAC at 20 mM/L for 1 or 2 h prior to the infection of CVB3, and then cells were infected with CVB3 and grown in the medium without the supplementation of NAC for 24 h. As shown in Figure 3A, massive production of ROS was seen in the cells infected with CVB3. The generation of ROS was markedly reduced in the CVB3-infected cells with NAC pretreatment (Figure 3A and B), while the synthesis of viral 3D^pol^ was not suppressed (Figure 3C). These data indicate that the antioxidant property of NAC does not contribute to its antiviral activity. Moreover, these data also suggest that NAC does not block virus entry in the course of CVB3 infection, since CVB3 was efficiently replicated in the cells pretreated with NAC (Figure 3C).

**Figure 3.**
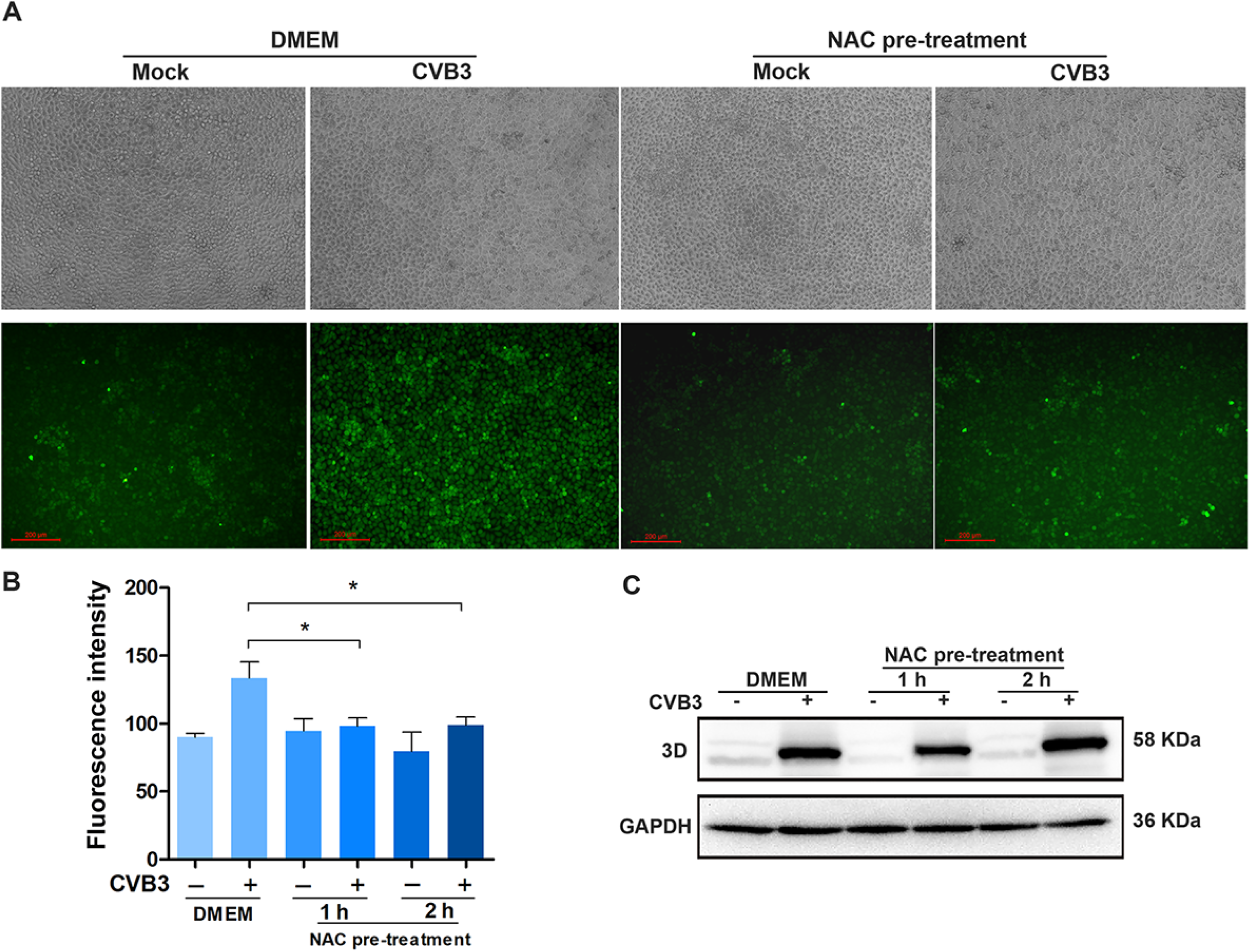
The antiviral effect of NAC is independent of its antioxidant property. HeLa cells were pretreated with NAC at 20 mM for 1 h and then infected or mock-infected with CVB3 at MOI of 1 for 24 h. (A) The generation of ROS was viewed in fluorescence microscope. (B) The relative fluorescence intensity was quantified. (C) Cells were pre-treated with 20 mM NAC for 1 or 2 h and then infected with CVB3 for 24 h. Cells were collected and cell lysates were subjected to Western blot analysis. Experiments were repeated three times. *n* = 3. \**P* < 0.05.

### NAC directly inhibits the activation of caspase-1

Viruses are obligate intracellular pathogens which utilize cellular machinery to synthesize macromolecules and to achieve efficient replication. Thus, we asked whether NAC regulates cellular components which are essential to viral replication. Our previous study demonstrated that selective caspase-1 inhibitor not only alleviated the inflammation caused by CVB3 infection, but also showed antiviral effect ^39^, suggesting the essential role of caspase-1 in CVB3 infection. In the present study, we observed the reduced production of pro-inflammatory cytokines including TNF-α, IL-6, IL-1β, and IL-18 during the infection of CVB3 in both mouse myocardium (Figure 1E) and HeLa cells (Figure S1A) treated with NAC. Given that caspase-1 is required for the maturation of both IL-1β and IL-18, the reduced IL-1β and IL-18 implies that caspase-1, its precursor or active form, is down-regulated by NAC treatment.

To this end, we infected HeLa cells with CVB3 (at MOI =1) and determined caspase-1 level at various time points of post-infection. As shown in Figure 4A, dramatically increased level of caspase-1 p20, one of the active fragments of caspase-1, was seen in the cells at 12 h of post-infection. As we have demonstrated previously ^39^, procaspase-1 and NLRP3, the critical constituents of NLRP3 inflammasome, were also increased due to CVB3 infection (Figure 4A). These data show that CVB3 infection not only induces the activation of caspase-1, but also increases the level of procaspase-1.

**Figure 4.**
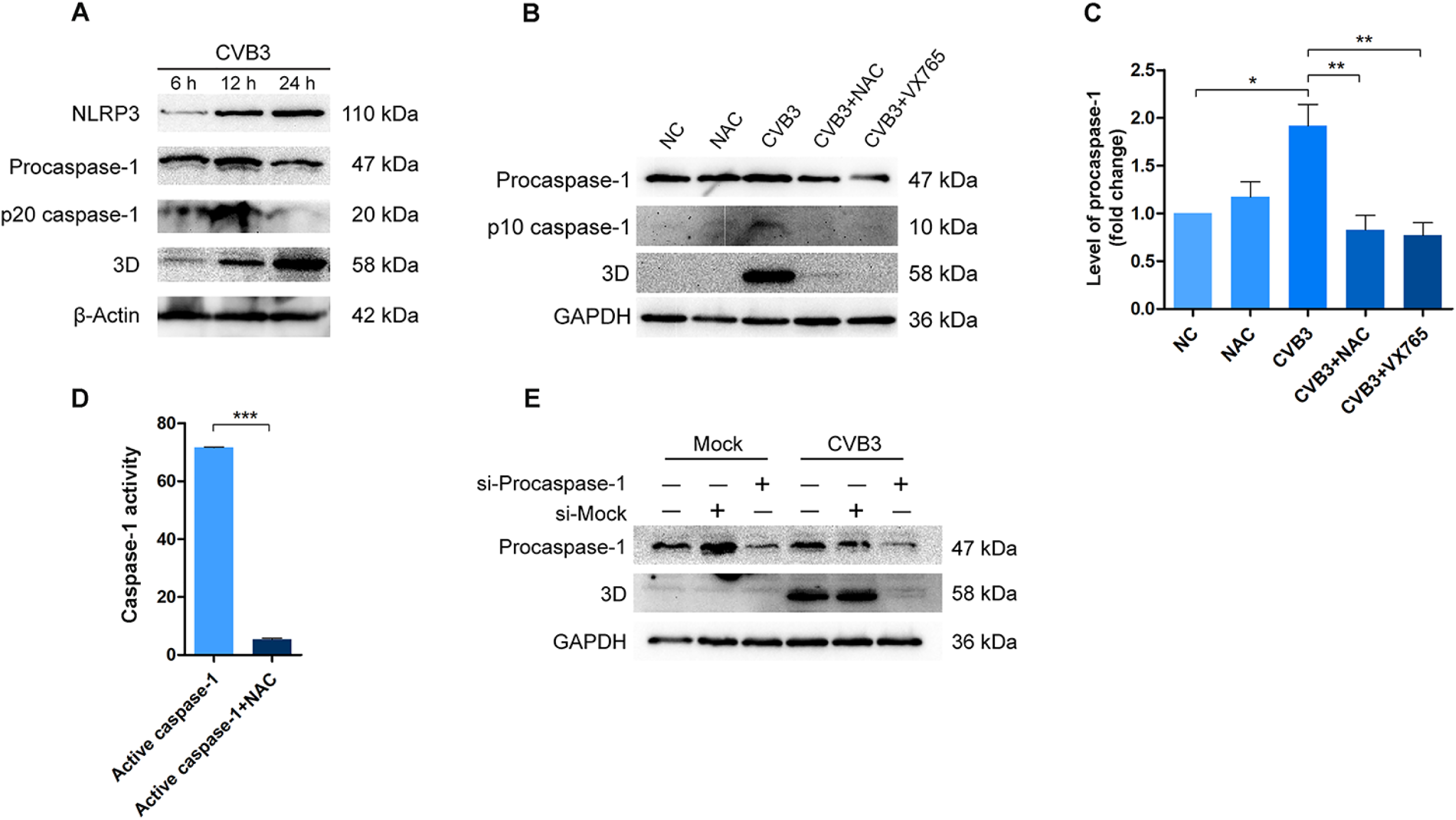
NAC inhibits caspase-1 activation. (A) HeLa Cells were infected with CVB3 at MOI of 1. Cells were collected at various time points of post-infection and analyzed by Western blotting. (B and C) HeLa cells were infected with CVB3 (MOI = 1) and treated with NAC or VX765 for 12 h. Cells were collected and cell lysates were analyzed by Western blotting. (D) 10 units of the recombinant active caspase-1 was incubated with 50 nM NAC at 37 °C for 2 h in the buffer provided by the Caspase-1 Activity Assay Kit. Caspase-1 activity was determined by a microplate reader at 405 nm. (F) HeLa cells were transfected with siRNA of procaspase-1 for 36 h. Control cells were transfected with scramble siRNA (si-Mock). Cells were then infected with CVB3 for 12 h. Cell lysates were analyzed by Western blotting. Experiments were repeated three times. *n* = 3. \**P* < 0.05; * \**P* < 0.01; * * \**P* < 0.001.

Although the role played by caspase-1 in CVB3 replication remains to be further studied, our data pointed the possibility that the anti-CVB3 effect of NAC might be related to its impact on caspase-1 activation. To address the question if NAC influences the activity of caspase-1 in the context of CVB3 infection, HeLa cells were infected with CVB3 and treated with NAC or caspase-1 inhibitor (VX765) for 12 h. The caspase-1 precursor and its active form (p10) were determined. CVB3 infection significantly increased both the precursor and active caspase-1 p10 (Figure 4B and C), while the treatment of NAC or caspase-1 inhibitor clearly reduced procaspase-1 as well as the active caspase-1 (p10) (Figure 4B and C). Importantly, the diminished active caspase-1 coincided with the disappearance of viral 3D^pol^ (Figure 4C). These data clearly show that NAC down-regulates caspase-1, and this action may contribute to the anti-CVB3 effect of NAC.

We next tried to answer the question how NAC exerts inhibition on caspase-1 activation. NAC is a thiol-containing chemical and the pro-drug of cysteine which provides sulfhydryl during its reactions ^54^. Thus, we postulated that the NAC’s inhibition on caspase-1 activity might be associated with the chemical character of this molecule, which allows it to form disulfide bonds with the cysteine residues of caspase-1 ^55^. To address this possibility, we performed *in vitro* study by testing the impact of NAC on the recombinant active caspase-1 (ab39901, Abcam; Cambridge, MA). 10 units of the recombinant caspase-1 was incubated with 50 nM of NAC at 37°C for 2 h, and caspase-1 activity was measured. As shown in Figure 4D, NAC treatment remarkably reduced the activity of caspase-1, indicating that NAC directly inhibits the activity of caspase-1.

To provide evidence that caspase-1 plays a role in CVB3 infection, we performed knockdown study using the siRNA of procaspase-1. HeLa cells were transfected with siRNA of procaspase-1 for 36 h and infected with CVB3 for 12 h. We found that the synthesis of 3D^pol^ of CVB3 was almost completely blocked in the cells with the knockdown of procaspase-1, indicating that caspase-1 is required for CVB3 replication.

Collectively, the above data show that the anti-CVB3 mechanism of NAC relies, at least partly on inhibiting caspase-1 activation, which is required for CVB3 replication.

### NAC promotes the degradation of procaspase-1 through ubiquitin-proteasome pathway

Even though the mechanism by which NAC inhibits caspase-1 activation has been studied (Figure 4C-E), we still need to answer the question how NAC down-regulates procaspase-1. To this end, we first determined the protein levels of procaspase-1 in the cells treated with MG132 or infected with CVB3 in the presence or absence of NAC. As shown in Figure 5, MG132 treatment obviously increased the levels of both procaspase-1 and procaspase-3 (Figure 5A-C), indicating that procaspases, at least for procaspase-1 and procaspase-3, are degraded through UPS. CVB3 infection also increased the levels of the precursors of both caspase-1 and caspase-3 (Figure 5A-C). Importantly, NAC treatment down-regulated the level of procaspase-1 in the cells with either MG132 treatment or CVB3 infection, suggesting that NAC promotes the degradation of procaspase-1.

**Figure 5.**
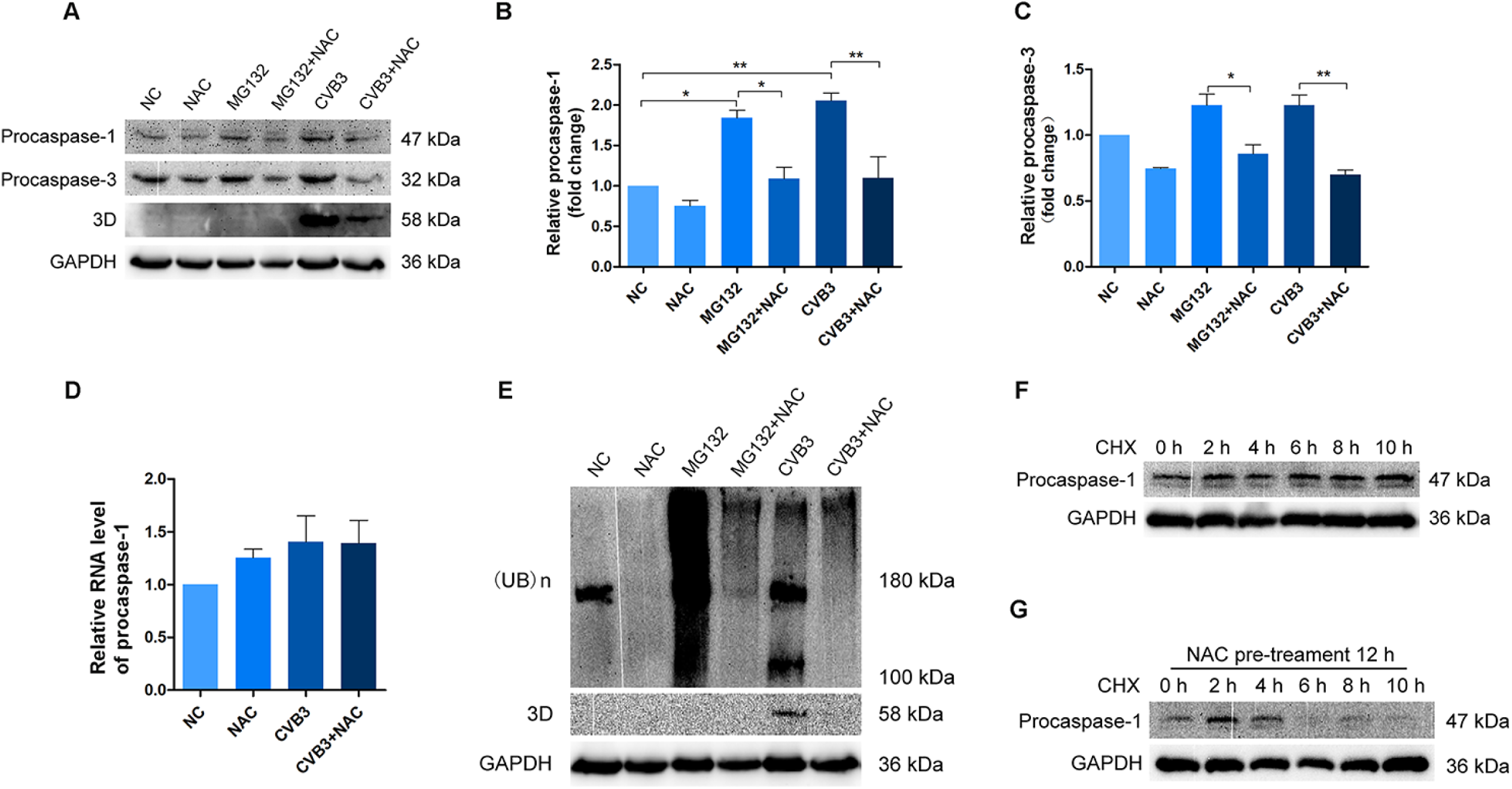
NAC promotes the degradation of procaspase-1. (A to C) HeLa cells were treated with MG132 or infected with CVB3 (MOI =1) for 12 h in the presence or absence of 20 mM NAC. Cells were collected and cell lysates were analyzed by Western blotting. The quantitative change of procaspase-1 and procaspase-3 were calculated (B and C). (D) HeLa cells were infected with CVB3 (MOI =1) in the presence or absence of 20 mM NAC for 12 h. Total RNAs were extracted and the abundance of procaspase-1 mRNA was determined by RT-qPCR. (E) HeLa cells were treated with MG132 or infected with CVB3 for 12 h (as described in A). Cell lysates were prepared and subjected to Western blot analysis. (F) HeLa cells were treated with 20 ng/ml cycloheximide (CHX). Cells were collected at various time points after CHX treatment and analyzed by Western blotting. (G) HeLa cells were pre-treated with 20 mM NAC for 12 and cultured in the medium containing CHX. Cell lysates were prepared at various time points of CHX treatment and analyzed by Western blotting. Experiments were repeated three times. *n* = 3. \**P* < 0.05; * \**P* < 0.01; * * \**P* < 0.001. (UB)n: polyubiquitinated proteins.

To further evaluate whether the expression of procaspase-1 is changed at transcriptional level, HeLa cells were infected with CVB3 in the presence of NAC for 12 h, and mRNA level of procaspase-1 was determined by RT-qPCR. As shown in Figure 5D, the abundance of procaspase-1 mRNA was not changed in the cells infected with CVB3 with or without the treatment of NAC, indicating that the transcription or stability of procaspase-1 mRNA is not interfered. These data also suggest that the protein degradation mechanism might play a role for the decline of procaspase-1 with NAC treatment.

26S proteasome is the protein assembly that mediates the degradation of polyubiquitin-conjugated proteins outside of lysosome. To show whether NAC regulates protein degradation through UPS, HeLa cells were treated with MG132 or infected with CVB3 for 12 h in the presence or absence of NAC. Ubiquitin-conjugated proteins were determined. As shown in Figure 5E, treatment of MG132 led to the vast accumulation of ubiquitin-conjugated proteins, while the polyubiquitinated proteins almost completely disappeared with NAC treatment. CVB3 infection also induced the accumulation of ubiquitinated proteins, which were distinct in molecular weight from the ubiquitin-conjugated proteins induced by MG132. Similarly, NAC also significantly reduced the ubiquitinated proteins induced by CVB3 infection. Taken together the results above (Figure 5 A to D), these data suggest that NAC promotes the degradation of procaspase-1 through regulating UPS.

To confirm that NAC promotes the degradation of procaspase-1, HeLa cells were treated with cycloheximide (CHX) (Jinpin Chemical, Shanghai, China), the inhibitor of protein synthesis. The amount of procaspase-1 was determined at various time points after CHX treatment. As shown in Figure 5F and G, within 10 h of CHX treatment, the amount of procaspase-1 has no obvious change. In contrast, in the cells pre-treated with NAC for 12 h, procaspase-1 disappeared at 6 h of CHX treatment, indicating that NAC accelerates the degradation of procaspase-1.

### NAC inhibits viral proteases

The enterovirus proteases of 2A^pro^ and 3C^pro^, which are essential for processing viral polyprotein during viral replication, are chymotrypsin-like cysteine proteases ^5^. Motivated by our finding that NAC inhibits the activity of caspase-1, we asked the question if NAC also shows inhibitory effect on viral proteases, since viral proteases, like caspase-1, are also cysteine proteases ^56^. In addition to processing viral polyproteins, 2A^pro^ and 3C^pro^ target multiple cellular proteins including eIF4G ^13^ and TAR DNA binding protein (TDP-43) ^57^. The cleavage of eIF4G contributes to the translational shutoff of the host cells during picornavirus infection ^13^, while the cleavage and degradation of TDP-43 by CVB3 has been shown to modulate viral pathogenesis ^57^.

To show the impact of NAC on the activity of viral proteases 2A^pro^ and 3C^pro^, HeLa cells were transfected with the construct pEGFP-2A or pEGFP-3C for 24 h. Cells were cultured for another 24 h in the presence of NAC. The proteolytic cleavage of eIF4G (Figure 6A) and TDP-43 (Figure 6B) were determined by Western Blotting. In the cells expressing EGFP-2A, eIF4G was cleaved and an extra fragment of about 130 kDa was observed, while the addition of NAC blocked the cleavage of eIF4G (Figure 4A). Similarly, the cleavage of TDP43 was observed in the cells expressing EGFP-3C, which generated a fragment in 35 kDa. NAC treatment for the cells expressing EGFP-3C blocked the cleavage of TDP-43 (Figure 6B). These results show that NAC suppresses the activity of viral 2A^pro^ and 3C^pro^.

**Figure 6.**
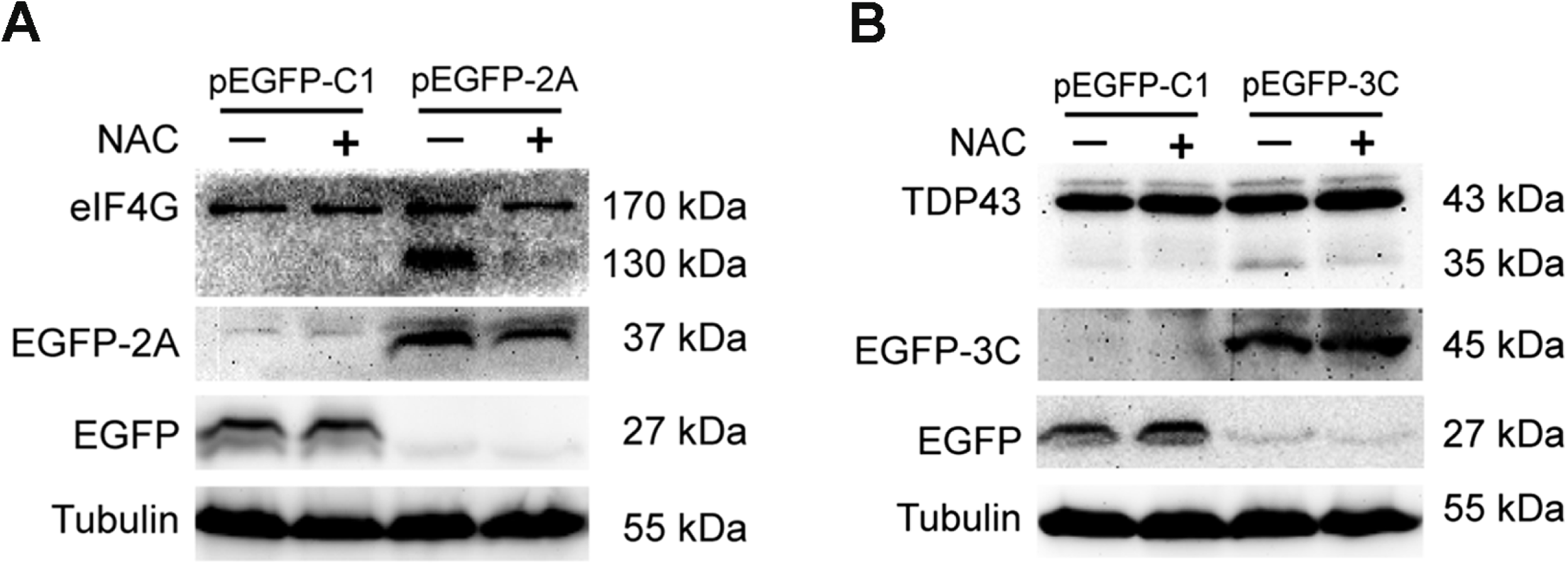
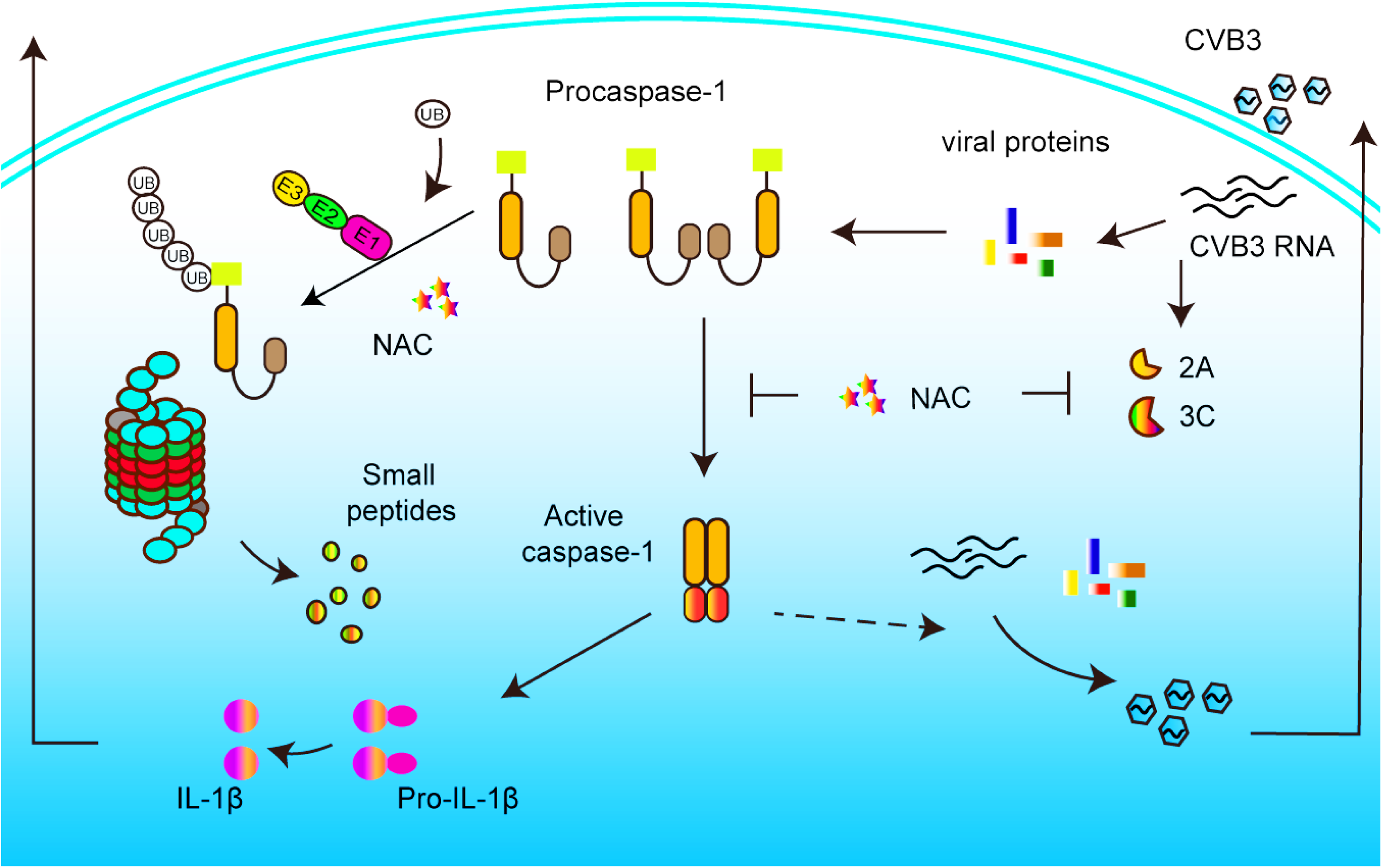
NAC inhibits the activity of viral proteases. (A) HeLa cells were transfected with pEGFP-2A for 24 h. Control cells were transfected with pEGFP-C1. Cell lysates were prepared and analyzed by Western blotting. (B) HeLa cells were transfected with pEGFP-3C for 24 h. Control cells were transfected with pEGFP-C1. Cell lysates were prepared and analyzed by Western blotting. Experiments were repeated three times.

## Discussion

The infection of CVB3 can cause severe and life-threatening viral myocarditis, which may progress to dilated cardiomyopathy ^9^. Until now, no effective antiviral medications are available for the treatment of CVB3 infection ^58^. The present study demonstrated that NAC, an antioxidant and the prodrug of cysteine, exhibits antiviral and anti-inflammatory effect against CVB3 infection both *in vitro* and *in vivo*. The antiviral mechanism of NAC is related to its inhibiton on caspase-1 and viral proteases. Based on this study, we propose that NAC could potentially be a safe and effective therapeutic option for myocarditis caused by CVB3 infection.

Myocarditis is defined as the inflammation of the cardiac muscle. The histological presentations of viral myocarditis caused by CVB3 infection include the necrosis of cardiomyocytes and the infiltration of lymphocytes or monocytes ^7, 59^. Cardiac inflammation caused by CVB3 infection is characterized with increased production of pro-inflammatory cytokines and the activation of nuclear factor kappa B (NF-kB) ^60^. Because of the lack of specific antiviral medicine, supportive care is still the primary treatment to CVB3-induced myocarditis ^58^.

NAC, as a water-soluble and thiol-containing antioxidant, is the established antidote for the treatment paracetamol overdose ^61^. It is recommended as the therapeutic option for conditions characterized by the generation of ROS ^62^. The -SH group of NAC confers it with the character of reducing disulfide bonds. Thus, NAC is widely used as mucolytic medicine to reduce the viscosity of the mucus generated during respiratory diseases such as COPD, cystic fibrosis, and chronic bronchitis ^63, 64^. Except as antioxidant, NAC also possesses anti-inflammatory activity through inhibiting the activation of NF-κB ^65^. The application of NAC for diabetic patients with dialysis significantly suppressed the activation of NF-κB ^48^.

The inflammatory character of viral myocarditis and the anti-inflammation activity of NAC motivated us to use NAC to combat the devastating inflammatory response induced by CVB3 infection. We were also encouraged by the report that blocking IL-1β-mediated signaling pathway led to significantly inhibited inflammation, viral replication, and cardiac changes ^66^. Thus, we used NAC to treat mice with viral myocarditis induced by CVB3 infection. We initially anticipated that NAC would alleviate myocardial inflammation and improve the overall condition of the infected mice. Unexpectedly, we found that, in addition to its anti-inflammation and cardio-protective effect, NAC also showed remarkable inhibition on CVB3 replication. To understand its antiviral mechanism, we started by determining the activity of caspase-1, since our previous study showed that CVB3 replication was inhibited by caspase-1 inhibitor ^39^. Indeed, we found that NAC not only inhibited caspase-1 activation, but also reduced the level of procaspase-1. We further demonstrated that the reduced procaspase-1 was due to its increased degradation through ubiquitin-proteasome pathway, as a result of NAC treatment. We also determined the impact of NAC on the activity of viral 2A^pro^ and 3C^pro^, because these proteases, similar to caspase-1, are also cysteine proteases. Our data show that in the presence of NAC, viral proteases lost their proteolytic activity towards cellular proteins eIF4G and TDP-43, respectively. Collectively, this study demonstrated that NAC exerts antiviral and cardio-protective effect for CVB3-induced myocarditis. The antiviral mechanism of NAC is associated with its inhibition on caspase-1 as well as viral proteases.

Obviously, our results contradict to the study by Si et al ^67^, in which no antiviral effect of NAC was observed against CVB3 infection in cultured cells. This difference is likely caused by the intrinsic reactivity of NAC, in which cysteine is oxidized to cystine at the contact of air in neutral or alkaline solution (Sigma-Aldrich, https://www.sigmaaldrich.com/catalog/product/sigma/a9165). Therefore, the concentration of NAC is gradually decreasing in working solution. This could be the reason that NAC lost its inhibitory effect on CVB3 replication. To maintain the precise concentration of NAC, we stored NAC at -80°C and prepared fresh working solution for each application.

For the mouse model of viral myocarditis, we show that when NAC was given at relative low dosage (15 mg/kg body weight, twice a day), significant cardio-protective and antiviral effect were achieved. According to the clinical application of NAC for the treatment of acetaminophen overdose or gentamycin toxicity, intravenous administration of NAC at 1200 mg twice a day is well tolerated ^62, 68-70^. Therefore, our results suggest that NAC could be a safe option for the treatment of CVB3 infection. However, further laboratory and clinical studies are needed.

In this study, we showed that when 20 mM of NAC was used in cell culture, maximum antiviral effect was obtained, while the cell viability was largely maintained. It is worth to note that the half-life of NAC is typically short ^71^. According to an *in vitro* study, only 18% free NAC was retained after incubating in serum containing medium for 24 h ^72^. It is believed that NAC is rapidly metabolized due to its incorporation on to proteins. In spite of its short half-life, we observed its antiviral effect at 24 h after the addition of NAC in cell culture. The antiviral and anti-inflammatory effect were also confirmed in mouse model of viral myocarditis. These data suggest that the antiviral mechanism of NAC may involve its influence on cellular and viral components, leading to the inhibition of inflammation and viral replication.

Similar to other picornaviruses, CVB3 infection induces oxidative stress ^51, 67^. Evidence also shows that ROS is harnessed by certain plant RNA virus to facilitate viral replication ^53^. Therefore, we evaluated whether the antioxidant property of NAC plays a role in its antiviral mechanism. Unlike the report about EV71 ^73^, in which the reduction of ROS suppressed the production of viral progeny, we demonstrated that the antiviral effect of NAC is independent of its antioxidant function.

The anti-inflammation effect of NAC inspired us to evaluate its influence on caspase-1, since we previously showed that caspase-1 inhibitor suppressed CVB3 replication ^39^. Here we demonstrated that NAC inhibits the activity of caspase-1 and down-regulates the level of procaspase-1 through regulating UPS. NAC is the synthetic precursor of cysteine. Both NAC and cysteine contain free sulfhydryl group which enable these molecules to function as nucleophilic and antioxidant. They can also form disulfide bonds with thiol-containing molecules such as the cysteine residues of proteins ^74^. In this study, we found that the activity of caspase-1 was directly inhibited by NAC (Figure 4F). This is likely the result of the reaction between NAC and the cysteine residue in the catalytic site of caspase-1, leading to the formation of disulfide bond and the conformational change of caspase-1. Consequently, caspase-1 is inactivated. This point of view is supported by the fact that the majority of caspase-1 inhibitors covalently interacting with the active site cysteine residue of the enzyme ^75, 76^. For example, VX-765, the inhibitor of caspase-1, functions through reversible covalent modification of the catalytic cysteine residue of the enzyme ^75^. Similarly, NAC’s inhibition on the proteases of CVB3 may also involves the same mechanism, because viral proteases also contain cysteine residue in their catalytic sites ^56^. Studies on hepatitis A virus (HAV) and human rhinovirus (HRV), which are important pathogens in the picornavirus family, showed that thiol-reactive groups such as iodoacetamides and beta-lactones can be used to inhibit the activity of 3C^pro^ of HAV and HRV ^77^.

We further demonstrated that UPS is involved in the antiviral mechanism of NAC. UPS, the important protein degradation machinery, participates in almost all cellular activities, and it plays crucial roles in maintaining the proteostasis of mammalian cells ^18^. Evidence shows that UPS is exploited by CVB3 ^17, 78^, while the precise role of UPS in CVB3 replication remains unknown. Moreover, how NAC would influence UPS in the context of CVB3 infection is unclear. In this study, we found that the protein levels of both procaspase-1 and procaspase-3 were increased by the treatment of MG132 (Figure 5A-C), the proteasome inhibitor, indicating that procaspases, at least for procaspase-1 and procaspase-3, are degraded through UPS. Importantly, NAC treatment blocked the accumulation of caspase-1 induced by either MG132 or CVB3 infection, and the decline of procaspase-1 was coincided with the significantly reduced production of viral 3D^pol^ (Figure 5A-C). We further show that the reduction of procaspase-1 is not due to the suppressed gene expression, because the mRNA level of procaspase-1 was not changed in the cells infected with CVB3 or treated with NAC (Figure 5D). Instead, we demonstrated that NAC regulates UPS and promotes the degradation of procaspase-1 (Figure 5E-G). Similarly, the degradation of p53, a cell cycle regulatory protein which is degraded through UPS, was also accelerated by the treatment of NAC (supplementary Figure S2).

How exactly can NAC regulate UPS remains unknown. Although our data showed that NAC accelerated the degradation of procaspase-1, we have no direct evidence that demonstrates the influence of NAC on the process of ubiquitination or proteasomal degradation. Our data suggest that NAC accelerates the degradation of the precursors of pan-caspases. In consistent to this postulation, a study has shown that E3 ligase TRAF2 exerts negative effect on the activity of caspase-8 ^79^. A previous study suggested that NAC improves the process of protein folding, which results in the disappearance of ubiquitinated proteins, while NAC did not subvert the inhibition of MG132 on proteasome ^80^. Nonetheless, if protein degradation were not involved in the decline of procaspase-1 in the presence of NAC, there would be no accelerated reduction of this protein when cells were treated with CHX, the inhibitor of protein synthesis (Figure 5F and G).

There is still one question which remains unanswered. If the degradation of procaspase-1, which would lead to the further inhibition of caspase-1, is not favorable for CVB3 replication, the suppressed degradation of procaspase-1 would facilitate viral replication. In contrast, MG132, the proteasome inhibitor which blocks the degradation of procaspase-1, also has anti-CVB3 effect ^81^. These data imply that capase-1 and UPS may exert their impact on CVB3 replication through distinct mechanisms. Nonetheless, the role of UPS in CVB3 infection needs to be clarified.

The limitation of this study is that we have not demonstrated the role of caspase-1 in CVB3 infection, although we believe that the activation of caspase-1 is crucial for CVB3 replication, since viral replication was almost blocked when procaspase-1 was knocked down. Furthermore, although NAC has been used in clinical practice for several decades as the prodrug of cysteine and the precursor of glutathione, the mechanisms underlying its therapeutic activity towards various conditions are still unclear. Evidence has shown that NAC influences multiple signaling pathways of the cell such NF-κB ^82^, p38 MAPK ^83^, JNK ^84^, and nitric oxide ^85^. Thus, our study certainly cannot exclude the possibility that critical cellular pathways might also contribute to the antiviral effect of NAC.

In conclusion, this study demonstrated that NAC exerts antiviral and anti-inflammatory effect against CVB3 infection. NAC treatment clearly alleviated the myocarditis and suppressed CVB3 replication. The antiviral mechanism of NAC is related with its inhibition on caspase-1 and viral proteases (Figure 7). Thus, NAC might be a safe therapeutic option for the treatment of myocarditis caused by CVB3 infection.

## Materials and Methods

### Ethical approval

All protocols were approved by the Ethics Committee of Harbin Medical University. All experimental procedures were conducted in accordance with the regulation on the use and care of laboratory animals for research.

### Mice

Newborn Balb/c mice were purchased from the Laboratory Animal Center of Harbin Medical University (Harbin, China). Mice were housed in specific pathogen-free environment at the temperature of 25 ± 1 °C and 40-50% humidity. Mice were allowed to access food and water ad libitum. To generate animal myocarditis model, newborn mice (n = 6-9 mice per group) were infected with 10^6^ TCID_50_ of CVB3 by intraperitoneal injection once at day 3-5 after birth. Control mice were injected with phosphate buffered saline (PBS) (Biosharp, Hefei, China). Mice were monitored and euthanized at day 5 of post-infection (p.i.). Mouse ventricles were collected and subjected to histological examination, RNA and protein extraction, and ELISA. Total of 96 new-born mice were used. Body weight change of each mouse was calculated according to the formula: (body weight – body weight of previous day) /5 day.

### Cell culture

HeLa cells were kindly provided by the Department of Medical Genetics of Harbin Medical University. Cells were grown in a humidified incubator with 5% CO_2_ at 37 °C. Cells were maintained in Dulbecco’s Modified Eagle Medium (DMEM) (Thermo Fisher, Shanghai, China) supplemented with 10% (v/v) fetal bovine serum (FBS) (Bioindustry, Israel), penicillin (100U/L), and streptomycin (100 U/L). Cells were provided with fresh medium every 2-3 day3. Cells infected with viruses were grown in the medium containing 2%FBS.

### Virus

CVB3 woodruff was kindly provided by the Scrips Institute (San Diego, USA) as previously described ^86^. Virus was amplified in HeLa cells and stored at -80°C. Virus stock was subjected to 3 times of freeze-thaw cycle and centrifuged 5 min at 1000 rpm to collect to the supernatant. Virus was titrated. Virus stock was passaged no more than three times to exclude possible cell adaptation. The 50% tissue culture infective dose (TCID_50_) of CVB3 used in this study was 10^7^/100 µL. HeLa cells at the confluency of 70% were infected with CVB3 at 1 of multiplicity of infection (MOI). Cells were allowed to absorb viruses for 1 h, and then the supernatant was removed and replaced with fresh media.

### NAC treatment

NAC (A9165; Sigma-Aldrich, St. Louis, MO) was dissolved in PBS. Aliquoted solution of NAC was stored at -80°C and used only once to avoid oxidation. HeLa cells infected or mock-infected with CVB3 were cultured in the medium containing 20 mM NAC for various time periods. For *in vivo* experiments, mice were infected or mock-infected with 10^3^ TCID50 CVB3. 15 mg/kg (body weight) of NAC was given intraperitoneally to mice twice a day beginning at 12 h of p.i.. Mice were euthanized at the end of day 5 of p.i..

### Transfection

Plasmids expressing the non-structural protein 2A or 3C of CVB3 (designated as pEGFP-2A and pEGFP-3C, respectively) and the control plasmid pEGFP-C1 were constructed as described previously ^12^. These constructs were amplified in the DH5 strain of E.coli (Takara, Dalian, China). Cells were seeded in 6-well plates at the concentration of 5 × 10^4^ cells/well and cultured for 24 h before transfection. To transfect cells, 500μL of DMEM containing 4μg plasmid mixed with 10μL Lipofectmine 2000 (Thermo Fisher) was added to each well of the 6-well culture plate when cells grew to 60-70% confluency. Transfection with interference RNA (siRNA) of procaspase-1 was performed similarly. After transfection, cells were allowed to grow continuously for 24 h in fresh culture medium. Cells were harvested and subjected to the analysis of RT-PCR and Western blot.

### RNA extraction and real-time quantitative PCR (RT-qPCR)

Cells were cultured in 6-well plate to 90% confluency and total RNA was extracted by Trizol (Life Technologies, Carlsbad, CA) and RT-qPCR was performed following the instruction of the manufacturer. Briefly, 1 μg of total RNA was reverse transcribed in a reaction of 10 µL with 4μL of 5×TransScript All-in-One SuperMix (Transgen, Beijing, China). Quantitative PCR was performed by adding 1 μL of cDNA, 0.4 μM of forward and reverse primers, 10 μL of 2×TransStart Top Green qPCR SuperMix (Transgen, Beijing, China), and RNase-free water to make 20 µL of total reaction. PCR reaction was carried out in LightCycler 96 (Roche) for 45 cycles of denaturation at 94 °C for 5 s,annealing at 55 °C for 15 s,and extension at 72°C for 1 min. The relative RNA amount was calculated with the 2^-ΔΔCT^ threshold cycle (CT) method and normalized to the amount of GAPDH ^87^. All reactions were carried out on triplicate. Primers were synthesized by Genewiz (Suzhou, China) and the sequences of the primers are follows: for CVB3, forward primer 5′-GCACACACCCTCAAACCAGA-3′ and reverse primer 5′-ATGAAACACGGACACCCAAAG-3′ ; for GAPDH, forward primer 5′-TGCACCACCAACTGCTTAGC-3′ and reverse primer 5′-GGCATGGACTGTGGTCATGAG-3′.

### Western blot

To extract proteins from cultured cells, cells grown in 6-well plate to 90% confluency were treated with 100 μL RIPA (Beyotime, Wuhan, China) containing protease inhibitor PMSF (v/v at 100:1) (Beyotime) for 20 min on ice. Cells were scraped from the culture plate and centrifuged at 4°C for 20 min to collect the supernatant. To extract proteins from tissue, mouse heart was collected and atrium was removed. Mouse ventricles were washed in cold PBS and homogenized in RIPA containing protease inhibitor PMSF for 5 min at ice. The homogenates were centrifuged at 12, 000 r/min for 20 min in 4°C. Supernatant was collected and stored at -80°C.

Protein concentration was determined by BCA protein assay kit (Pierce, Rockford, USA). Protein lysates were separated in 10% SDS-PAGE gel (Transgen, Beijing, China) by Mini-PROTEAN tetra Cell system (BioRad) and transferred to PVDF membrane. PVDF membrane was washed, blocked in skimmed milk for 1 h and incubated with primary antibody for 2 h at room temperature. Blots were developed with ECL (Boster, Wuhan, China) and imaged by FluorChem R system (Protein Simple). Polyclonal antibodies against EGFP, eIF4G, capase-1, caspase-3, TDP-43, β-actin, α-tubulin, and GAPDH were obtained from Proteintech (Wuhan, China). Anti-enterovirus VP1 monoclonal antibody was obtained from DAKO (M 7064; Dako, Denmark). Anti-3D of CVB3 rabbit polyclonal antibody was prepared in our laboratory.

### Enzyme-linked immunosorbent assay (ELISA)

The supernatant of cell culture was collected. IL-1β, IL-6, IL-18, and TNF-α were determined by the enzyme-linked immunosorbent assay (ELISA) following the instructions of the provider (Elabscience, Wuhan, China). After color development, the optical density at 450 nm was determined by microplate reader Epoch 2 (BioTek, Winooski, VT). To determine mouse IL-1β, IL-6, IL-18, and TNF-α, mouse hearts were collected and homogenized. The supernatant was collected for ELISA.

### Histopathology

Mouse hearts were collected and the ventricles were fixed and embedded in paraffin. Cardiac tissues were sectioned and subjected to HE staining as described previously. Tissue sections were analyzed separately by two specialists in the Department of Pathology of Harbin Medical University.

### Cell viability

HeLa cells were seeded in 96-well plates and treated with NAC (Sigma-Aldrich) at various concentrations for 24 h. Cell viability was determined using Cell Counting Kit-8 (CCK-8) (Beyotime, Wuhan, China) in accordance with the protocol of the provider. In brief, after the incubation period, the culture medium was removed and fresh medium with 20 μl of 5 mg/ml CCK-8 reagent was added. Cells were incubated for another 2 h until the formation of formazan crystals. Cells were washed with PBS for three times, and formazan crystals were solubilized with 150 μl of DMSO. Cell viability was determined using a microplate reader Epoch2 (BioTek) at 570 nm and normalized to control cells. Positive control was also provided by the manufacturer.

### Measurement of ROS production

To measure ROS production, cells were pre-treated with NAC at 20 mM for 1 h and infected with CVB3 (MOI =1) for 24 h. Cells were incubated with 10 µM DCF-DA (Beyotime, Wuhan, China), a fluorogenic dye that measures hydroxyl, peroxyl and other ROS activities within the cell, for 30 min. Cells were then washed twice with ice-cold PBS followed the observation of by Leica DM2000 fluorescence microscope (Germany) at the wavelength of 485/520 nm (absorption/emission). Fluorescence intensity was calculated with the ImageJ.

### Caspase-1 activity assay

Caspase-1 activity was measured using Caspase 1 Activity Assay Kit (Solarbio, Beijing, China) according to the instruction of the manufacturer. 10 unit of active recombinant human caspase-1 protein (ab39901, Abcam; Cambridge, MA) and 50 nM NAC were added to the reaction buffer, which was incubated at 37 °C for 2 h. A standard curve was prepared. The optical density was read on a microplate reader (Epoch2, BioTek) at 405 nm.

### Statistical analysis

*In vitro* experiments were repeated at least four times. Animal experiments were repeated three times with 6 to 9 mice per treatment group. Quantitative data are presented as mean ± SD. Student *t* test was used to compare the results. *P* value of less than 0.05 is considered as statistically significant. Data were analyzed by Graphpad Prism 8.

## Acknowledgements

This study was supported by the National Natural Science Foundation of China (Grant 81672007 to Wenran Zhao, 81571999 and 81871652 to Zhaohua Zhong, and 81772188 to Yan Wang). We thank the technical support by the Center of Northern Translational Medicine and the Institute of Wu Lian-Tai of Harbin Medical University. We thank Mr Hongbo Chui and Ms Yuehui Zhao (Laboratory Center of Microbiology, Harbin Medical University, Harbin, China) for their excellent technical support for the care of laboratory mice.

## Author Contributions

Yao Wang, S Zhao, and Y Chen performed the majority of the experiments. Q Cong and X Feng were responsible for animal care. J Zhang and Y Dong assisted in the collection of data. Z Zhong and W Zhao conceived and designed the study. Z Zhong, W Zhao, and Yao Wang analyzed the data. W Zhao and Z Zhong wrote the manuscript. The remaining authors provided substantial help for the implementation of this study. All authors read and agreed to the manuscript.

## Conflict of interest

There is no conflict of interests in this work.

## References

1. Huber, S.A. Viral Myocarditis and Dilated Cardiomyopathy: Etiology and Pathogenesis. Curr Pharm Des 22, 408–426 (2016).

2. Yajima, T. & Knowlton, K.U. Viral myocarditis: from the perspective of the virus. Circulation 119, 2615–2624 (2009).

3. Knowlton, K.U. & Lim, B.K. Viral myocarditis: is infection of the heart required? J Am Coll Cardiol 53, 1227–1228 (2009).

4. Uhl, T.L. Viral myocarditis in children. Crit Care Nurse 28, 42–63; quiz 64 (2008).

5. Garmaroudi, F.S. et al. Coxsackievirus B3 replication and pathogenesis. Future Microbiol 10, 629–653 (2015).

6. Bowles, N.E. et al. Analysis of the coxsackievirus B-adenovirus receptor gene in patients with myocarditis or dilated cardiomyopathy. Mol Genet Metab 77, 257–259 (2002).

7. Doan, D., Rungta, S., Vikraman, N. & Rosman, H. Fulminant Coxsackie B myocarditis mimicking acute coronary artery occlusion. Tex Heart Inst J 37, 500–501 (2010).

8. Persichino, J., Garrison, R., Krishnan, R. & Sutjita, M. Effusive-constrictive pericarditis, hepatitis, and pancreatitis in a patient with possible coxsackievirus B infection: a case report. BMC Infect Dis 16, 375 (2016).

9. Menahem, S. Viral myocarditis and dilated cardiomyopathy in early childhood. Br Heart J 58, 420–421 (1987).

10. Yajima, T. Viral myocarditis: potential defense mechanisms within the cardiomyocyte against virus infection. Future Microbiol 6, 551–566 (2011).

11. Feng, Q. et al. Enterovirus 2Apro targets MDA5 and MAVS in infected cells. J Virol 88, 3369–3378 (2014).

12. Wu, S. et al. Protease 2A induces stress granule formation during coxsackievirus B3 and enterovirus 71 infections. Virol J 11, 192 (2014).

13. Chau, D.H. et al. Coxsackievirus B3 proteases 2A and 3C induce apoptotic cell death through mitochondrial injury and cleavage of eIF4GI but not DAP5/p97/NAT1. Apoptosis 12, 513–524 (2007).

14. Pestova, T.V., Shatsky, I.N. & Hellen, C.U. Functional dissection of eukaryotic initiation factor 4F: the 4A subunit and the central domain of the 4G subunit are sufficient to mediate internal entry of 43S preinitiation complexes. Mol Cell Biol 16, 6870–6878 (1996).

15. Joachims, M., Van Breugel, P.C. & Lloyd, R.E. Cleavage of poly(A)-binding protein by enterovirus proteases concurrent with inhibition of translation in vitro. J Virol 73, 718–727 (1999).

16. Gao, G. et al. Proteasome inhibition attenuates coxsackievirus-induced myocardial damage in mice. Am J Physiol Heart Circ Physiol 295, H401–408 (2008).

17. Si, X. et al. Dysregulation of the ubiquitin-proteasome system by curcumin suppresses coxsackievirus B3 replication. J Virol 81, 3142–3150 (2007).

18. Nandi, D., Tahiliani, P., Kumar, A. & Chandu, D. The ubiquitin-proteasome system. J Biosci 31, 137–155 (2006).

19. Pant, V. & Lozano, G. Limiting the power of p53 through the ubiquitin proteasome pathway. Genes Dev 28, 1739–1751 (2014).

20. Urano, T. et al. p57(Kip2) is degraded through the proteasome in osteoblasts stimulated to proliferation by transforming growth factor beta1. J Biol Chem 274, 12197–12200 (1999).

21. Van Linthout, S. & Tschope, C. Viral myocarditis: a prime example for endomyocardial biopsy-guided diagnosis and therapy. Curr Opin Cardiol 33, 325–333 (2018).

22. Huber, S. Tumor necrosis factor-alpha promotes myocarditis in female mice infected with coxsackievirus B3 through upregulation of CD1d on hematopoietic cells. Viral Immunol 23, 79–86 (2010).

23. Lane, J.R., Neumann, D.A., Lafond-Walker, A., Herskowitz, A. & Rose, N.R. Role of IL-1 and tumor necrosis factor in coxsackie virus-induced autoimmune myocarditis. J Immunol 151, 1682–1690 (1993).

24. Huber, S.A., Gauntt, C.J. & Sakkinen, P. Enteroviruses and myocarditis: viral pathogenesis through replication, cytokine induction, and immunopathogenicity. Adv Virus Res 51, 35–80 (1998).

25. Muller, I. et al. CX3CR1 knockout aggravates Coxsackievirus B3-induced myocarditis. PLoS One 12, e0182643 (2017).

26. Remels, A.H.V. et al. NF-kappaB-mediated metabolic remodelling in the inflamed heart in acute viral myocarditis. Biochim Biophys Acta Mol Basis Dis 1864, 2579–2589 (2018).

27. Keller, M., Ruegg, A., Werner, S. & Beer, H.D. Active caspase-1 is a regulator of unconventional protein secretion. Cell 132, 818–831 (2008).

28. Thornberry, N.A. et al. A novel heterodimeric cysteine protease is required for interleukin-1 beta processing in monocytes. Nature 356, 768–774 (1992).

29. Elliott, J.M., Rouge, L., Wiesmann, C. & Scheer, J.M. Crystal structure of procaspase-1 zymogen domain reveals insight into inflammatory caspase autoactivation. J Biol Chem 284, 6546–6553 (2009).

30. Swanson, K.V., Deng, M. & Ting, J.P. The NLRP3 inflammasome: molecular activation and regulation to therapeutics. Nat Rev Immunol (2019).

31. Shrivastava, G., Leon-Juarez, M., Garcia-Cordero, J., Meza-Sanchez, D.E. & Cedillo-Barron, L. Inflammasomes and its importance in viral infections. Immunol Res 64, 1101–1117 (2016).

32. Lupfer, C., Malik, A. & Kanneganti, T.D. Inflammasome control of viral infection. Curr Opin Virol 12, 38–46 (2015).

33. Yin, Y. et al. Early hyperlipidemia promotes endothelial activation via a caspase-1-sirtuin 1 pathway. Arterioscler Thromb Vasc Biol 35, 804–816 (2015).

34. Ferrer, L.M. et al. Caspase-1 Plays a Critical Role in Accelerating Chronic Kidney Disease-Promoted Neointimal Hyperplasia in the Carotid Artery. J Cardiovasc Transl Res 9, 135–144 (2016).

35. Kanneganti, T.D. Inflammatory Bowel Disease and the NLRP3 Inflammasome. N Engl J Med 377, 694–696 (2017).

36. Wang, Y. et al. Inflammasome Activation Triggers Caspase-1-Mediated Cleavage of cGAS to Regulate Responses to DNA Virus Infection. Immunity 46, 393–404 (2017).

37. Zheng, Y. et al. Zika virus elicits inflammation to evade antiviral response by cleaving cGAS via NS1-caspase-1 axis. EMBO J 37 (2018).

38. Cai, R. et al. Caspase-1 Activity in CD4 T Cells Is Downregulated Following Antiretroviral Therapy for HIV-1 Infection. AIDS Res Hum Retroviruses 33, 164–171 (2017).

39. Wang, Y. et al. Pyroptosis induced by enterovirus 71 and coxsackievirus B3 infection affects viral replication and host response. Sci Rep 8, 2887 (2018).

40. Roth, L., Adler, M., Jain, T. & Bempong, D. Monographs for medicines on WHO’s Model List of Essential Medicines. Bull World Health Organ 96, 378–385 (2018).

41. Saritas, A. et al. N-Acetyl cysteine and erdosteine treatment in acetaminophen-induced liver damage. Toxicol Ind Health 30, 670–678 (2014).

42. Alsalim, W. & Fadel, M. Towards evidence based emergency medicine: best BETs from the Manchester Royal Infirmary. Oral methionine compared with intravenous n-acetyl cysteine for paracetamol overdose. Emerg Med J 20, 366–367 (2003).

43. Hoffer, B.J. et al. Repositioning drugs for traumatic brain injury - N-acetyl cysteine and Phenserine. J Biomed Sci 24, 71 (2017).

44. Harrison, P.M., Wendon, J.A., Gimson, A.E., Alexander, G.J. & Williams, R. Improvement by acetylcysteine of hemodynamics and oxygen transport in fulminant hepatic failure. N Engl J Med 324, 1852–1857 (1991).

45. He, G. et al. N-Acetylcysteine for Preventing of Acute Kidney Injury in Chronic Kidney Disease Patients Undergoing Cardiac Surgery: A Metaanalysis. Heart Surg Forum 21, E513–E521 (2018).

46. Tung, W.H., Hsieh, H.L., Lee, I.T. & Yang, C.M. Enterovirus 71 induces integrin beta1/EGFR-Rac1-dependent oxidative stress in SK-N-SH cells: role of HO-1/CO in viral replication. J Cell Physiol 226, 3316–3329 (2011).

47. Sreekanth, G.P. et al. Drug repurposing of N-acetyl cysteine as antiviral against dengue virus infection. Antiviral Res 166, 42–55 (2019).

48. Amore, A. et al. N-Acetylcysteine in hemodialysis diabetic patients resets the activation of NF-kB in lymphomonocytes to normal values. J Nephrol 26, 778–786 (2013).

49. Lee, S.I. & Kang, K.S. N-acetylcysteine modulates lipopolysaccharide-induced intestinal dysfunction. Sci Rep 9, 1004 (2019).

50. Zhao, Y., Wang, M., Li, Y. & Dong, W. Andrographolide attenuates viral myocarditis through interactions with the IL-10/STAT3 and P13K/AKT/NF-kappabeta signaling pathways. Exp Ther Med 16, 2138–2143 (2018).

51. Wang, Y., Gao, B. & Xiong, S. Involvement of NLRP3 inflammasome in CVB3-induced viral myocarditis. Am J Physiol Heart Circ Physiol 307, H1438–1447 (2014).

52. Zhitkovich, A. N-Acetylcysteine: Antioxidant, Aldehyde Scavenger, and More. Chem Res Toxicol (2019).

53. Hyodo, K., Hashimoto, K., Kuchitsu, K., Suzuki, N. & Okuno, T. Harnessing host ROS-generating machinery for the robust genome replication of a plant RNA virus. Proc Natl Acad Sci U S A 114, E1282–E1290 (2017).

54. Kashfi, K. & Olson, K.R. Biology and therapeutic potential of hydrogen sulfide and hydrogen sulfide-releasing chimeras. Biochem Pharmacol 85, 689–703 (2013).

55. Kim, H.J., Ha, S., Lee, H.Y. & Lee, K.J. ROSics: chemistry and proteomics of cysteine modifications in redox biology. Mass Spectrom Rev 34, 184–208 (2015).

56. Malcolm, B.A. The picornaviral 3C proteinases: cysteine nucleophiles in serine proteinase folds. Protein Sci 4, 1439–1445 (1995).

57. Fung, G. et al. Cytoplasmic translocation, aggregation, and cleavage of TDP-43 by enteroviral proteases modulate viral pathogenesis. Cell Death Differ 22, 2087–2097 (2015).

58. Pollack, A., Kontorovich, A.R., Fuster, V. & Dec, G.W. Viral myocarditis--diagnosis, treatment options, and current controversies. Nat Rev Cardiol 12, 670–680 (2015).

59. Zhai, X. et al. Coxsackievirus B3 Induces Autophagic Response in Cardiac Myocytes in vivo. Biochemistry (Mosc) 80, 1001–1009 (2015).

60. Zhou, L., He, X., Gao, B. & Xiong, S. Inhibition of Histone Deacetylase Activity Aggravates Coxsackievirus B3-Induced Myocarditis by Promoting Viral Replication and Myocardial Apoptosis. J Virol 89, 10512–10523 (2015).

61. Mokhtari, V., Afsharian, P., Shahhoseini, M., Kalantar, S.M. & Moini, A. A Review on Various Uses of N-Acetyl Cysteine. Cell J 19, 11–17 (2017).

62. Atkuri, K.R., Mantovani, J.J., Herzenberg, L.A. & Herzenberg, L.A. N-Acetylcysteine--a safe antidote for cysteine/glutathione deficiency. Curr Opin Pharmacol 7, 355–359 (2007).

63. Santus, P. et al. Oxidative stress and respiratory system: pharmacological and clinical reappraisal of N-acetylcysteine. COPD 11, 705–717 (2014).

64. Biswas, S., Hwang, J.W., Kirkham, P.A. & Rahman, I. Pharmacological and dietary antioxidant therapies for chronic obstructive pulmonary disease. Curr Med Chem 20, 1496–1530 (2013).

65. Haddad, J.J. Antioxidant and prooxidant mechanisms in the regulation of redox(y)-sensitive transcription factors. Cell Signal 14, 879–897 (2002).

66. Kraft, L., Erdenesukh, T., Sauter, M., Tschope, C. & Klingel, K. Blocking the IL-1beta signalling pathway prevents chronic viral myocarditis and cardiac remodeling. Basic Res Cardiol 114, 11 (2019).

67. Si, X. et al. Pyrrolidine dithiocarbamate reduces coxsackievirus B3 replication through inhibition of the ubiquitin-proteasome pathway. J Virol 79, 8014–8023 (2005).

68. Khayyat, A., Tobwala, S., Hart, M. & Ercal, N. N-acetylcysteine amide, a promising antidote for acetaminophen toxicity. Toxicol Lett 241, 133–142 (2016).

69. Senanayake, M.P., Jayamanne, M.D. & Kankananarachchi, I. N-acetylcysteine in children with acute liver failure complicating dengue viral infection. Ceylon Med J 58, 80–82 (2013).

70. Lim, G. & Lee, J.H. N-acetylcysteine in children with dengue-associated liver failure: a case report. J Trop Pediatr 58, 409–413 (2012).

71. Borgstrom, L. & Kagedal, B. Dose dependent pharmacokinetics of N-acetylcysteine after oral dosing to man. Biopharm Drug Dispos 11, 131–136 (1990).

72. Holdiness, M.R. Clinical pharmacokinetics of N-acetylcysteine. Clin Pharmacokinet 20, 123–134 (1991).

73. Ho, H.Y. et al. Glucose-6-phosphate dehydrogenase deficiency enhances enterovirus 71 infection. J Gen Virol 89, 2080–2089 (2008).

74. Poole, L.B. The basics of thiols and cysteines in redox biology and chemistry. Free Radic Biol Med 80, 148–157 (2015).

75. Boxer, M.B. et al. A highly potent and selective caspase 1 inhibitor that utilizes a key 3-cyanopropanoic acid moiety. ChemMedChem 5, 730–738 (2010).

76. Romanowski, M.J., Scheer, J.M., O’Brien, T. & McDowell, R.S. Crystal structures of a ligand-free and malonate-bound human caspase-1: implications for the mechanism of substrate binding. Structure 12, 1361–1371 (2004).

77. Lall, M.S., Jain, R.P. & Vederas, J.C. Inhibitors of 3C cysteine proteinases from Picornaviridae. Curr Top Med Chem 4, 1239–1253 (2004).

78. Si, X. et al. Ubiquitination is required for effective replication of coxsackievirus B3. PLoS One 3, e2585 (2008).

79. Jin, Z. et al. Cullin3-based polyubiquitination and p62-dependent aggregation of caspase-8 mediate extrinsic apoptosis signaling. Cell 137, 721–735 (2009).

80. Gleixner, A.M. et al. N-Acetyl-l-Cysteine Protects Astrocytes against Proteotoxicity without Recourse to Glutathione. Mol Pharmacol 92, 564–575 (2017).

81. Luo, H. et al. Proteasome inhibition reduces coxsackievirus B3 replication in murine cardiomyocytes. Am J Pathol 163, 381–385 (2003).

82. Wang, H.W. et al. N-acetylcysteine administration is associated with reduced activation of NF-kB and preserves lung dendritic cells function in a zymosan-induced generalized inflammation model. J Clin Immunol 33, 649–660 (2013).

83. Zhang, R., Wang, Y., Pan, L. & Tian, H. N-Acetylcysteine potentiates the haemodynamic-improving effect of sildenafil in a rabbit model of acute pulmonary thromboembolism via the p38 MAPK pathway. Clin Exp Pharmacol Physiol 46, 163–172 (2019).

84. Onyango, I.G., Tuttle, J.B. & Bennett, J.P., Jr. Activation of p38 and N-acetylcysteine-sensitive c-Jun NH2-terminal kinase signaling cascades is required for induction of apoptosis in Parkinson’s disease cybrids. Mol Cell Neurosci 28, 452–461 (2005).

85. Anfossi, G. et al. N-acetyl-L-cysteine exerts direct anti-aggregating effect on human platelets. Eur J Clin Invest 31, 452–461 (2001).

86. Tong, L. et al. MiR-10a* up-regulates coxsackievirus B3 biosynthesis by targeting the 3D-coding sequence. Nucleic Acids Res 41, 3760–3771 (2013).

87. Livak, K.J. & Schmittgen, T.D. Analysis of relative gene expression data using real-time quantitative PCR and the 2(-Delta Delta C(T)) Method. Methods 25, 402–408 (2001).

